## Supplementary Information for "Dissecting the effect of single- and co-infection of TB and COVID-19 pathogens on the sputum microbiome"

### Contents


**Definitions of key measures** **2**

**Network and interaction analysis** **3**

**Supplementary Tables** **5**

**Supplementary Figures** **8**

**Supplementary Files** **15**

### Definitions of key measures

This section gives the definitions of the different measures/metrics used in our study.

**Relative abundance (also denoted as proportion $p$):** Given a sample, the relative abundance is calculated by dividing the count $x_i$ of a particular taxon $i$ in the sample by the total count of all $s$ taxa within the sample.

$$p_i = \frac{x_i}{\sum_{j=1}^{s} x_j}$$

**CLR transformation:** Given a sample, the CLR transformation is applied by taking the logarithm of the ratio of the count $x_i$ of a particular taxon $i$ in the sample to the geometric mean of all $s$ taxa counts within the sample.

$$CLR_i = log \frac{x_i}{\left(\prod_{j=1}^{s} x_j\right)^{\frac{1}{s}}}$$

The same transformation can also be applied to a vector of non-count data for a sample, such as the vector of pathway activity scores of a sample estimated by PICRUSt2.

**Alpha ($\alpha$) diversity:** Different $\alpha$-diversity measures were used to assess the distribution of taxa within a community [1]. We have used the following $\alpha$-diversity measures in this study.

1. Shannon (diversity) index: This index quantifies the uncertainty in the distribution of taxa within a sample and is given by:

   $$\text{Shannon index} = -\sum_{i=1}^{s} p_i \ln p_i$$

   Here, $s$ is the total number of taxa and $p_i$ is the relative abundance of taxa $i$ in a sample as calculated above.

2. Observed richness/taxa: This measure represents the total number of distinct taxa observed in a sample (i.e., the number of taxa $i$ for which $p_i > 0$).

3. Simpson (diversity) index: This measure gives the probability that two randomly selected individuals from a community will have the same taxa. Its formula is given by:

   $$\text{Simpson index} = 1 - \sum_{i=1}^{s} p_i^2$$

**Bray-Curtis dissimilarity:** It is a statistical measure used to quantify the dissimilarity in taxa composition between two samples from the same group.

$$\text{Bray-Curtis dissimilarity(A, B)} = \frac{\sum_{i=1}^{s} |a_i - b_i|}{\sum_{i=1}^{s} (a_i + b_i)}$$

Here, $a_i$ and $b_i$ are the abundances (read counts) of taxa $i$ in samples A and B respectively.

**ICC:** It is a statistical measure used to assess the degree of agreement or consistency between two or more different measurements or raters and is calculated based on the estimates of population variances. We have used the ICC absolute agreement formula which states that "Each target is rated by a different judge and the judges are selected at random" [2]. In our analysis, repeated samples represent the target, and Shannon diversity serves as the rating. The number of repeats correspond to the number of judges, representing sequencing performed across different batches.

$$ICC(1,1) = \rho_{1,1} = \frac{\sigma_r^2}{\sigma_r^2 + \sigma_w^2} \tag{1}$$

Here, $\sigma_r^2$ denotes the variance between targets, and $\sigma_w^2$ the variance within targets (across sequencing batches).

**Dispersion analysis:** It is used to assess the degree of variability (or spread) of microbial composition within or between data groups. We have used betadisper function of vegan package in R to evaluate the homogeneity of group dispersions between each pair of disease groups [3, 4]. This function calculates the distances of each sample from its group centroid in an ordination space based on the Bray–Curtis dissimilarity matrix, providing a measure of variability within the group. ANOVA was then used to test significant differences in variabilities between the groups.

### Network and interaction analysis

We compared the species-species association network of TB group with that of the TBCOVID group to identify species pairs that are differentially associated between the two groups. The networks were constructed using the SpiecEasi method, which considers associations as conditional dependence relationships between taxa after accounting for all other taxa in the community. On visually examining the two networks (Fig. S6), we observed a few shared associations such as *Rothia mucilaginosa*-*Granulicatella adiacens* and *Dialister invisus*-*Dialister pneumosimtes*, and several associations that were present in the TB but not in the TBCOVID network. But a statistical test of differential association (Fisher's z-test at significance level 0.05 after multiple testing correction) did not reveal any significant rewiring of the network between the groups. We also used the interaction analysis framework [5] developed by our lab to quantify the interaction effects in the presence of co-infections and no significant interactions were detected at the genus and species levels.

#### Supplementary Tables

Table. S1: Summary of the clinical characteristics of TB and TBCOVID groups generated using the tableone [6] package in Python. Here, BPL indicates Below Poverty Line. IP refers to the Intensive Phase of TB treatment (initial phase, typically 2 months), while CP denotes the Continuation Phase (follow-up phase, typically 4–6 months). Pos-smokeless indicates positive for smokeless tobacco use. RIF, INH, FQ, SLI, and SLI (eis) indicate resistance to Rifampicin, Isoniazid, Fluoroquinolones, Second-Line Injectable drugs (SLI), and Kanamycin due to enhanced intracellular survival (eis) gene-associated resistance, respectively. P-values were calculated using default tests in tableone, viz., Chi-squared test for categorical variables and ANOVA for continuous variables.

| | | Grouped by COVID Result | | | |
|---|---|---|---|---|---|
| | | Overall | Negative | Positive | p-value |
| n | | 48 | 24 | 24 | |
| Age (in years), mean (SD) | | 52.2 (11.4) | 52.5 (10.4) | 52.0 (12.5) | 0.861 |
| | range (min - max) | 25 - 80 | 36 - 75 | 25 - 80 | |
| Gender, n (%) | female | 7 (14.6) | 3 (12.5) | 4 (16.7) | 1.000 |
| | male | 41 (85.4) | 21 (87.5) | 20 (83.3) | |
| Socio Economic Status, n (%) | BPL | 38 (79.2) | 20 (83.3) | 18 (75.0) | 0.544 |
| | Unknown | 10 (20.8) | 4 (16.7) | 6 (25.0) | |
| New or Previously treated, n (%) | New | 48 (100.0) | 24 (100.0) | 24 (100.0) | 1.000 |
| Weight (in kilograms), mean (SD) | | 46.2 (10.3) | 45.5 (10.8) | 46.9 (9.9) | 0.658 |
| | range (min - max) | 27 - 69 | 32 - 69 | 27 - 65 | |
| Height (in centimeters), mean (SD) | | 158.8 (6.7) | 158.8 (6.9) | 158.7 (6.8) | 0.970 |
| | range (min - max) | 144 - 176 | 148 - 176 | 144 - 168 | |
| Treatment regimen, n (%) | Normal | 48 (100.0) | 24 (100.0) | 24 (100.0) | 1.000 |
| Treatment phase, n (%) | CP | 8 (16.7) | 3 (12.5) | 5 (20.8) | 0.701 |
| | IP | 40 (83.3) | 21 (87.5) | 19 (79.2) | |
| Treatment outcome, n (%) | Cured | 32 (66.7) | 16 (66.7) | 16 (66.7) | 0.347 |
| | Lost to follow up | 6 (12.5) | 4 (16.7) | 2 (8.3) | |
| | On Treatment | 5 (10.4) | 1 (4.2) | 4 (16.7) | |
| | Treatment Complete | 4 (8.3) | 3 (12.5) | 1 (4.2) | |
| | Died | 1 (2.1) | | 1 (4.2) | |
| HIV status, n (%) | Non Reactive / Negative | 48 (100.0) | 24 (100.0) | 24 (100.0) | 1.000 |
| Diabetes, n (%) | Diabetic | 27 (56.2) | 13 (54.2) | 14 (58.3) | 1.000 |
| | Non-diabetic | 21 (43.8) | 11 (45.8) | 10 (41.7) | |
| Tobacco-Smoking, n (%) | No | 31 (64.6) | 15 (62.5) | 16 (66.7) | 0.345 |
| | Unknown | 1 (2.1) | 1 (4.2) | | |
| | Yes | 14 (29.2) | 8 (33.3) | 6 (25.0) | |
| | Pos-smokeless | 2 (4.2) | | 2 (8.3) | |
| Alcohol, n (%) | No | 25 (52.1) | 11 (45.8) | 14 (58.3) | 0.563 |
| | Yes | 23 (47.9) | 13 (54.2) | 10 (41.7) | |
| Rifampicin, n (%) | Not Detected | 47 (97.9) | 24 (100.0) | 23 (95.8) | 1.000 |
| | - | 1 (2.1) | | 1 (4.2) | |
| Smear, n (%) | 1+positive | 19 (39.6) | 8 (33.3) | 11 (45.8) | 0.297 |
| | 2+positive | 15 (31.2) | 10 (41.7) | 5 (20.8) | |
| | 3+positive | 14 (29.2) | 6 (25.0) | 8 (33.3) | |
| RIF, n (%) | Not detected | 48 (100.0) | 24 (100.0) | 24 (100.0) | 1.000 |
| INH(InhA), n (%) | Not detected | 48 (100.0) | 24 (100.0) | 24 (100.0) | 1.000 |
| INH(KatG), n (%) | Not detected | 47 (97.9) | 24 (100.0) | 23 (95.8) | 1.000 |
| | Detected | 1 (2.1) | | 1 (4.2) | |
| FQ class resistance, n (%) | Not detected | 1 (100.0) | | 1 (100.0) | 1.000 |
| SLI, n (%) | Not detected | 1 (100.0) | | 1 (100.0) | 1.000 |
| SLI (eis), n (%) | Not detected | 1 (100.0) | | 1 (100.0) | 1.000 |
| COVID Result, n (%) | Negative | 24 (50.0) | 24 (100.0) | | $< 0.001$ |
| | Positive | 24 (50.0) | | 24 (100.0) | |

Table. S2: Overview of sample distribution across different sequencing batches for each group

| Group/Batch | Batch 1 | Batch 2 | Batch 3 | Batch 4 | Batch 5 | Total | Number of samples excluding repeat samples |
|---|---|---|---|---|---|---|---|
| **TB** | 4 | 2 | 12 | 8 | 1 | **27** | **24** |
| **COVID** | 2 | 4 | 1 | 1 | 4 | **12** | **10** |
| **TBCOVID** | 4 | 2 | 11 | 10 | 0 | **27** | **24** |
| **Control** | 2 | 4 | 1 | 1 | 18 | **26** | **24** |
| **Total** | **12** | **12** | **25** | **20** | **23** | **92** | **82** |

### Supplementary Figures

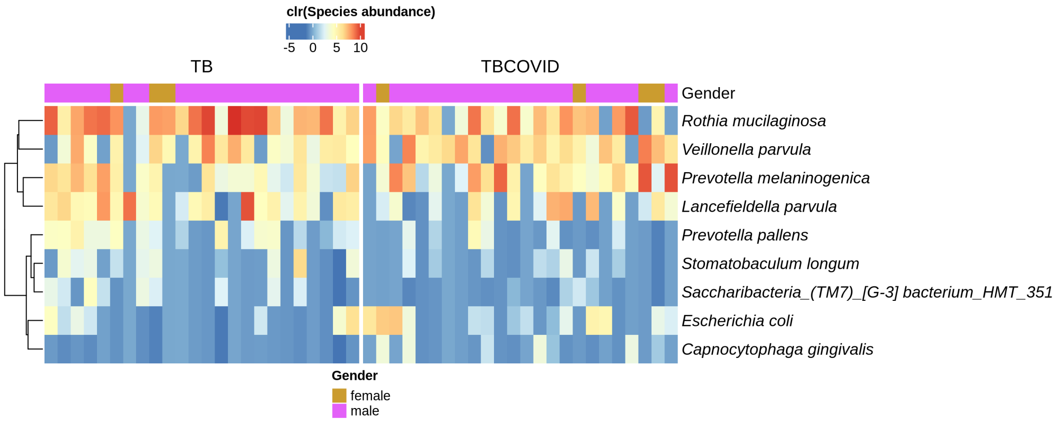


Fig. S1: **Heatmap of differentially abundant species:** The heatmap shows species found to be differentially abundant between the TB and TB-COVID groups (see Fig. 5 in the main text for details). Gender is indicated in the top annotation bar.

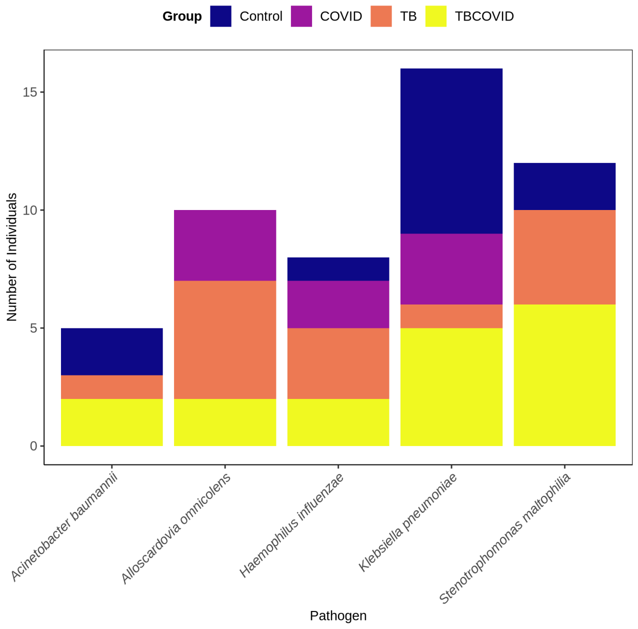


Fig. S2: **Pathogen analysis:** Distribution of respiratory pathogens across the four groups.

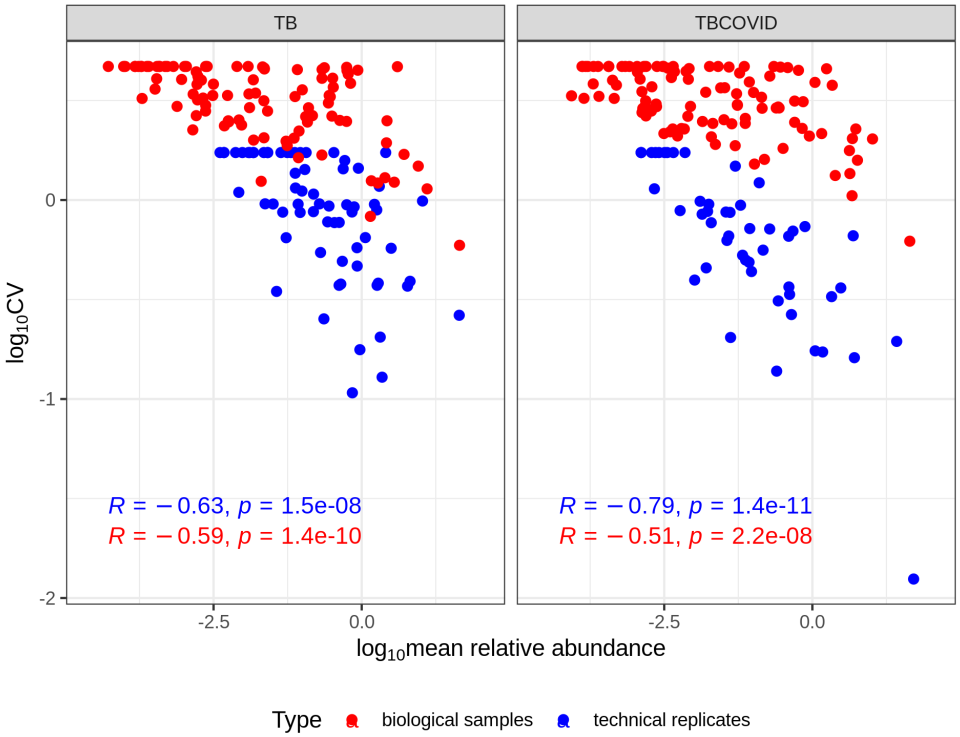


Fig. S3: **CV versus genus abundance across biological samples and technical replicates:** A sample with three technical replicates was used to assess technical variation. Samples without technical replicates were used to evaluate biological variation. Each point in the plot represents a genera. To expand a bit more, to assess technical variation, we subjected a TB sample to three repeat measurements – the CV and mean of the relative abundance of each genera across these three technical replicates is shown as a blue dot in the left panel. Similar plot is shown for a TBCOVID sample with three technical replicates in the right panel.

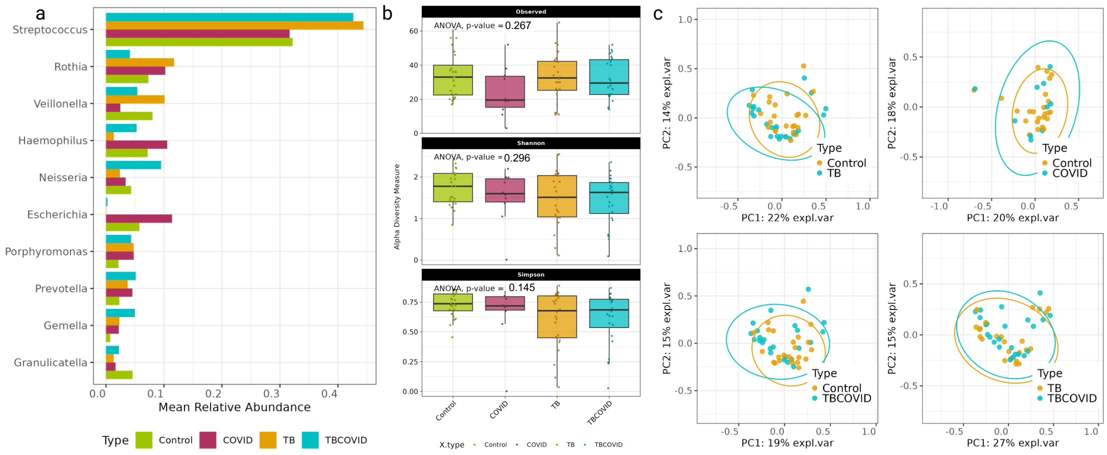


Fig. S4: **Characterization of genus diversity and composition in metagenomic sequencing data obtained from the four groups:** a) Visualization of the top 10 genera across the four groups, ranked based on their mean relative abundance. b) Boxplots of the alpha diversity measures (Observed, Shannon and Simpson) at genus level across the 4 groups. c) PCoA plots based on the Bray-curtis dissimilarity measure computed between the genus abundance profiles of samples.

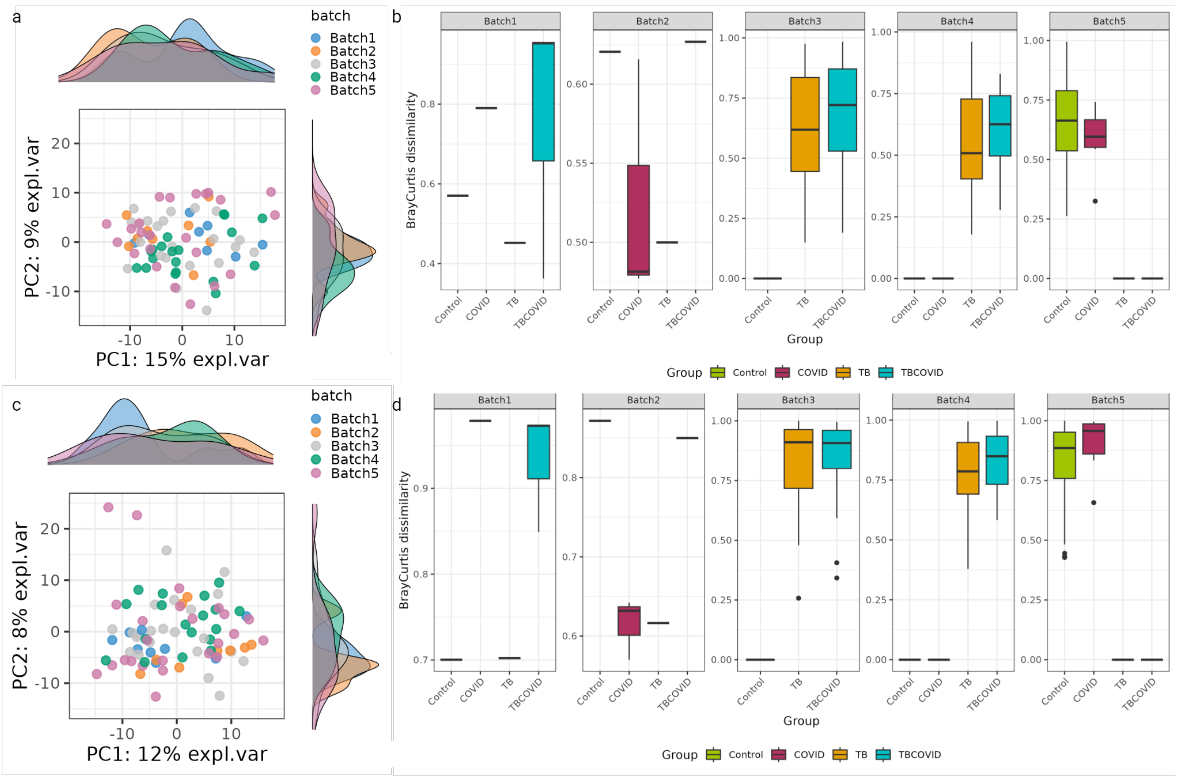


Fig. S5: **Batch effect analysis :** a) PCA plot of genus abundance across batches b) Bray-Curtis dissimilarity of samples within each group across batches based on genus abundance c) PCA plot of species abundance across batches d) Bray-Curtis dissimilarity of samples within each group across batches based on species abundance.

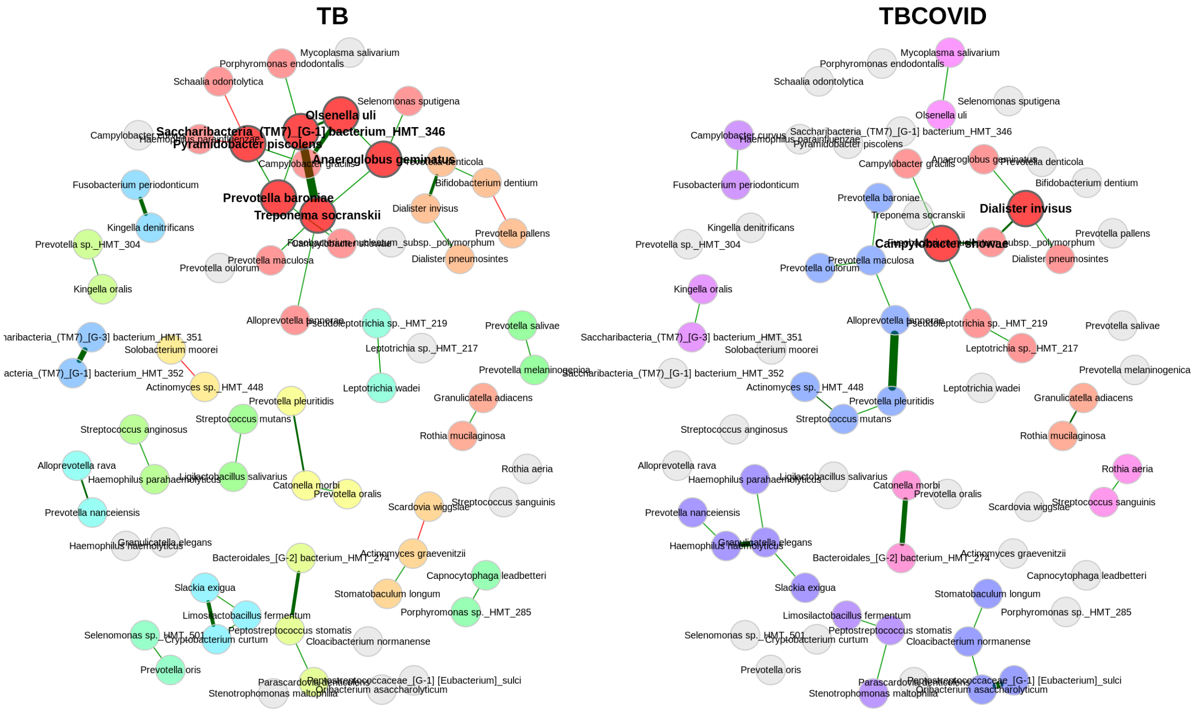


Fig. S6: **Network analysis results:** Association networks of TB and TBCOVID groups constructed using SpiecEasi method. The green and red colors of edges indicate positive and negative associations respectively. The thickness of edges corresponds to the strength of association, with thicker edges representing stronger associations. Only taxa present in at least 10% of samples within each group were used in the network construction.

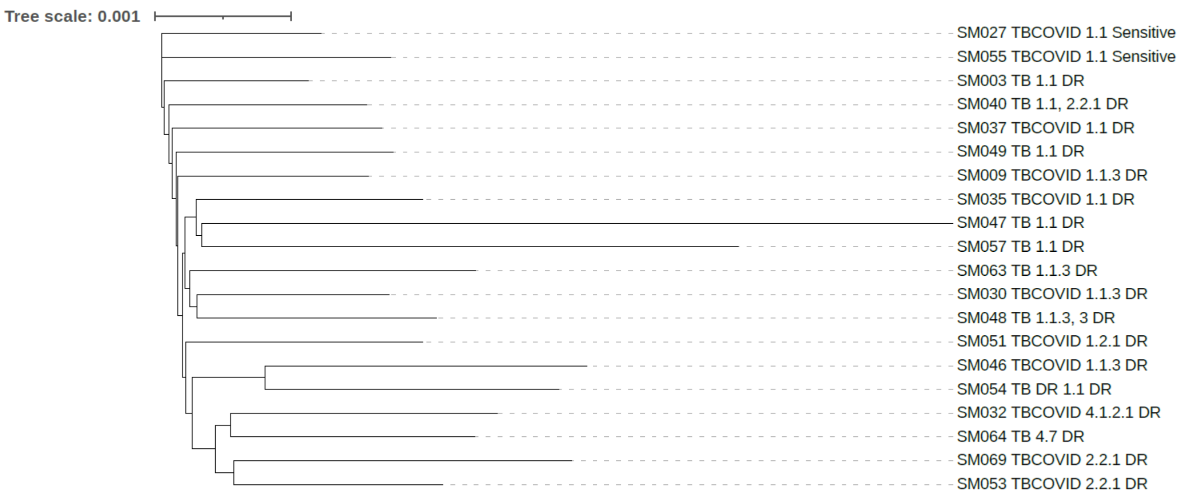


Fig. S7: **Genome analysis results:** Phylogenetic tree of the 9 TB and 11 TBCOVID samples constructed using the FastME/OneClick pipeline from the NGPhylogeny platform. The labels on the leaves indicate the group (TB/TB-COVID), lineages identified and drug resistance status (whether drug-resistant (DR) or sensitive). Two TB samples (SM040 and SM048) were identified as mixed infections containing two different strains of *M. tb*, with the major strain belonging to lineage 1.

#### Supplementary Files

Supplementary data/result files listed below are available at this link:
https://drive.google.com/drive/folders/1o7sikxfRSLLFTl3VZUGB8rAAxy4tlANZ?usp=drive_link.

Suppl File D1a: The covariates of all 461 TB individuals from whom sputum specimens were obtained.

Suppl File D1b: Sample identifiers (IDs) and the associated covariates of samples selected for sequencing across the four groups including replicates.

Suppl File D2: This file contains the raw and post-filtering read counts for each sample.

Suppl File D3: This file contains abundance data at the **genus**, **species**, and **pathway** levels. Genus and species read counts were generated using custom R scripts, while pathway abundances were obtained from PICRUSt2 analysis.

Suppl File D4: This file contains results from **genus**-level DA analysis for different comparisons of two groups (chosen from the four groups: TB-only, COVID-only, TBCOVID and Controls). The results of each pairwise comparison is in a separate tab in this xlsx file, and include the p-values and adjusted p-values (q-value) obtained by the three methods corncob, ANCOM-BC and LinDA for each **genus**. Note that corncob does not give log fold change (LFC) and only the LFC's for ANCOM-BC and corncob are included.

Suppl File D5: This file contains results from **species**-level DA analysis for different comparisons of two groups (chosen from the four groups: TB-only, COVID-only, TBCOVID and Controls). The results of each pairwise comparison is in a separate tab in this xlsx file, and the p-values and adjusted p-values (q-value) obtained by the three methods corncob, ANCOM-BC and LinDA for each **species**. Note that corncob does not give LFC and only the LFC's for ANCOM-BC and corncob are included.

Suppl File D6: This file contains results from **pathway**-level DA analysis for different comparisons of two groups (chosen from the four groups: TB-only, COVID-only, TBCOVID and Controls). The results of each pairwise comparison is in a separate tab in this xlsx file, and the p-values and adjusted p-values (q-value) obtained by the three methods corncob, ANCOM-BC and LinDA for each **pathway**. Note that corncob does not give LFC and only the LFC's for ANCOM-BC and corncob are included.
